# The Maine Coon Cat Harboring the *MYBPC3*-A31P Mutation: A Genotype-Stratified Phenotypic Characterization of Hypertrophic Cardiomyopathy

**DOI:** 10.64898/2026.08.13.744747

**Authors:** Xiaohe Shi, Ruoxuan Li, Zixuan Yang, Yue Wang, Junzhe Huang, Kai Liu, Jing Wang, Liwen Liu, Bo Wang

## Abstract

**Background:** Most animal models of HCM are mouse-based, but the thin interventricular septum in mice makes it difficult to clearly distinguish pathological hypertrophy, which introduces substantial errors and constrains basic HCM research. Cats develop HCM spontaneously, and the common *MYBPC3*-A31P variant in cats is homologous to human mutations in both genetics and pathology, with a larger body size that makes them suitable as large-animal models. This study examines how heterozygosity or homozygosity for the p.A31P mutation (c.91G>C) in the *MYBPC3* gene affects the phenotype and severity of HCM in affected cats, with the aim of establishing an ideal large-animal model for clinical risk stratification and precision diagnosis and treatment of human HCM.

**Methods:** Forty-nine Maine Coon cats were enrolled and stratified into homozygous mutant (HOM, n=8), heterozygous mutant (HET, n=26), and wild-type (WT, n=15) groups. All cats underwent echocardiography, blood pressure measurement, physiological assessment, hematological and biochemical analyses, and cross-species sequence conservation analysis.

**Results:** No significant differences in baseline characteristics including age and body weight were observed among groups (*P*>0.05). HOM cats exhibited significantly higher left ventricular outflow tract pressure gradients and greater basal septal thickness compared to WT cats (*P*<0.05), with HET cats showing intermediate values. Analysis of hematological and serum biochemical parameters revealed no evidence of systemic inflammation or hepatic injury. Sequence conservation analysis confirmed that the A31 residue is highly conserved across mammalian species.

**Conclusions:** This study provides a phenotypic characterization of Maine Coon cats carrying the *MYBPC3*-A31P mutation, revealing marked gene-dose effects on cardiac structure and function, with homozygous individuals exhibiting more severe phenotypic features. This model serves as a large-animal translational platform that not only clarifies genotype–phenotype correlations but also supports risk stratification and precision therapeutic strategies in human HCM. Its spontaneous nature and genetic homology to human disease make it particularly valuable for bridging preclinical findings to clinical application.

**RESEARCH PERSPECTIVE:** *What Is New?:*

- This study provides a genotype-stratified phenotypic characterization of *MYBPC3*-A31P mutation in Maine Coon cats, covering echocardiographic and laboratory parameters across homozygous, heterozygous, and wild-type groups.
- The results suggest a gene-dose effect, with homozygotes showing more pronounced cardiac changes, heterozygotes showing intermediate features, and wild-types remaining unaffected.
- All cats were raised under uniform environmental and dietary conditions, which may reduce the confounding effects of extrinsic factors compared with studies using cats from varied sources.

*What Question Should Be Addressed Next?:*

- Longitudinal follow-up of this cohort could help determine whether the observed genotype-related differences progress to clinical events over time.
- Protein-level and functional studies would be useful to explore the molecular mechanisms underlying the gene-dose-dependent phenotype. The utility of this feline model for evaluating genotype-specific therapeutic approaches remains to be examined.

## Introduction

Hypertrophic cardiomyopathy (HCM) is a genetic cardiac disease caused by mutations in genes encoding sarcomeric proteins. It is a leading cause of heart failure and sudden cardiac death in young adults^[1]^. Among humans, although significant progress has been made in understanding their genetic basis, there still lacks an ideal animal model. Most studies still rely on pharmacologically or surgically induced models—such as Angiotensin II infusion or transverse aortic constriction—which, while valuable for studying pressure overload, fail to recapitulate the molecular pathogenesis^[2]^. The most commonly used genetic models, transgenic mice, carry artificially introduced mutations that are not spontaneously occurring. Moreover, the murine heart presents significant anatomical limitations: interventricular septum thickness in mice is typically less than 1 mm^[3]^, making the distinction between hypertrophic and non-hypertrophic states technically challenging and often reliant on subtle statistical differences rather than unequivocal pathological thickening.

Cats, by contrast, develop HCM spontaneously with strong breed predisposition, showing high prevalence in Maine Coon, Ragdoll, and Persian cats^[4, 5]^. In Maine Coons, HCM is predominantly linked to an A31P mutation in the *MYBPC3* gene, which is inherited in an autosomal dominant pattern with incomplete penetrance and variable expressivity^[6, 7]^. This naturally occurring mutation is homologous to those found in human patients, positioning the Maine Coon as a promising translational model. Most existing studies in cats have focused on single echocardiographic parameters without integrating multi-dimensional phenotypic data across genetically confirmed groups^[8, 9]^. However, a well-characterized, genotype-stratified feline model with comprehensive phenotyping has been lacking. This gap limits the ability to fully understand genotype-phenotype correlations and hampers the model’s translational utility.

Therefore, by genotype-based screening of a Maine Coon cohort, we identified cats carrying the *MYBPC3*-A31P mutation and performed comprehensive phenotyping— including echocardiography, blood pressure, hematological and serum biochemical parameters—to systematically delineate clinical characteristics across wild-type, heterozygous, and homozygous genotypes. This naturally occurring feline model shares key genetic and pathological features with human HCM, thereby offering a valuable platform for exploring genotype–phenotype correlations and serving as a robust preclinical tool for translational research.

## Materials and Methods

### 2.1. Animals and Ethics

A total of forty-nine clinically healthy Maine Coon cats were enrolled. All cats were sourced from collaborative catteries with known HCM1 genotypic status. The HCM phenotype was defined according to established echocardiographic criteria, with an end-diastolic maximal left ventricular wall thickness of ≤5 mm considered normal, 5– 6 mm classified as equivocal, and ≥6 mm on at least one examination considered diagnostic for HCM, after excluding secondary causes of myocardial thickening such as systemic hypertension and hyperthyroidism^[10]^.This study was approved by the Institutional Animal Care and Use Committee of Yiji (Shanghai) Biotechnology Co., Ltd. (Approval No. YJ-2025-NA0015). All experimental procedures and animal welfare protocols complied with the relevant guidelines of the National Institutes of Health for the care and use of laboratory animals.

### 2.2. DNA sequencing and Genotyping

Blood samples (2-3 mL) were collected from each cat with minimal restraint and stored at −20°C until DNA extraction. Genomic DNA was extracted from 200 µL of whole blood using a commercial kit (DP348, Tiangen Biotech Co., Ltd., Beijing, China). Polymerase chain reaction (PCR) was performed using primers AB1P-E (5′-AGCAAGAAGCCAAGGTCAGT-3′) and AB1P-R (5′-CACTGGCGCTGATGTCAC-3′) under the following conditions: 95°C for 15 min; 35 cycles of 95°C for 30 s, 58°C for 30 s, and 72°C for 1 min; and a final extension at 72°C for 10 min. PCR products were purified and subjected to Sanger sequencing.

### 2.3. Clinical and Phenotypic Data Collection

All assessments were completed within two weeks by a fixed veterinary team to minimize inter-operator variability. Age, body weight, and sex were recorded. Echocardiographic examinations were performed using a Vivet300 system on cats in lateral recumbency. Measurements included left ventricular wall thickness, left atrial and aortic root dimensions, left ventricular ejection fraction (LVEF), and left ventricular outflow tract pressure gradient (LVOT-PG). Systolic blood pressure was measured non-invasively at the femoral artery. Heart rate and respiratory rate were determined by 15-second palpation and observation, respectively. Rectal temperature was measured digitally. Daily food and water consumption was quantified over 24 hours, with abnormal behaviors noted. The cat’s cooperation during restraint, coat condition, and skin status were also documented.

### 2.4. Statistical Analysis

All statistical analyses were conducted using SPSS software (Version 26.0). Continuous variables were first assessed for normality using the Shapiro-Wilk test and for homogeneity of variances using Levene’s test. For data that followed a normal distribution and exhibited homogeneity of variances, one-way analysis of variance (ANOVA) was used to compare differences among the three genotype groups, followed by post-hoc pairwise comparisons using the least significant difference (LSD) or Tukey’s test. For data that deviated from normal distribution or showed heterogeneity of variances, the *Kruskal-Wallis H* test was employed to assess overall differences among the groups. When a statistically significant difference was indicated by the *Kruskal-Wallis H* test (*P* < 0.05), post-hoc pairwise comparisons were conducted using the *Mann-Whitney U* test with *Bonferroni* correction for multiple comparisons. A two-tailed *P*-value of < 0.05 was considered statistically significant for all analyses.

## Results

### 3.1. Baseline Demographics and Group Characteristics

Based on the sequencing results, cats were classified into three genotypic groups: homozygous mutant (HOM, A31P/A31P, n=8), heterozygous mutant (HET, A31P/N, n=26), and wild-type (WT, N/N, n=15). Based on the predefined echocardiographic criteria (normal: end-diastolic maximal wall thickness ≤5 mm; equivocal: 5–6 mm; HCM: ≥6 mm), the phenotypic distribution of left ventricular wall thickness across the three groups was as follows. In the HOM (A31P/A31P) group (n=8), 5 cats (62.5%) exhibited a normal wall thickness and 3 cats (37.5%) met the diagnostic criteria for HCM. In the HET (A31P/N) group (n=26), 19 cats (73.1%) were classified as normal, 7 cats (26.9%) as equivocal, and none were diagnosed with HCM. In the WT (N/N) group (n=15), 14 cats (93.3%) were normal, 1 cat (6.7%) was equivocal, and no cat reached the HCM threshold.

The demographic profile of the 49 Maine Coon cats is summarized in Table 1. Maine Coon cats with different *MYBPC3* genotypes showed no statistically significant differences in baseline demographic and general physiological characteristics, including age, body length, weight, shoulder height, sex distribution, heart rate, body temperature, respiratory rate, water/food intake, and coat/skin condition (all *P* > 0.05). The systolic blood pressure (SBP) values for each group were as follows: 121.13 ± 7.90 mmHg in the HOM group, 128.00 ± 13.40 mmHg in the HET group, and 135.13 ± 13.45 mmHg in the WT group. The overall difference among the three groups did not reach statistical significance (*P* = 0.058).

**Table 1.** Baseline Demographics of the Study Population by MYBPC3 Genotype.

| Characteristic | HOM<br>(A31P/A31P)<br>(n=8) | HET<br>(A31P/N)<br>(n=26) | WT<br>(N/N)<br>(n=15) | *P*-value |
| --- | --- | --- | --- | --- |
| Age (years), Mean ± SD | 1.38 ± 0.63 | 1.14 ± 0.34 | 1.30 ± 0.24 | 0.215 |
| Body length (cm), Mean ± SD | 52.44 ± 6.32 | 52.89 ± 4.28 | 52.60 ± 5.17 | 0.957 |
| Weight (kg), Mean ± SD | 5.90 ± 1.53 | 5.71 ± 1.34 | 5.48 ± 1.60 | 0.934 |
| Shoulder height (cm), Mean ± SD | 31.62 ± 2.62 | 32.12 ± 2.79 | 32.07 ± 3.39 | 0.880 |
| Sex, Male n (%) | 6 (75.00%) | 18 (69.23%) | 11 (73.33%) | 0.924 |
| SBP (mmHg), Mean ± SD | 121.13 ± 7.90 | 128.00 ± 13.40 | 135.13 ± 13.45 | 0.058 |
| Heart rate (bpm), Mean ± SD | 158.00 ± 29.47 | 156.77 ± 15.12 | 169.33 ± 20.43 | 0.309 |
| Body Temp ($^{\circ}$C), Mean ± SD | 38.69 ± 0.33 | 38.75 ± 0.46 | 38.61 ± 0.52 | 0.344 |
| Respiratory rate (breaths/min) | 25.00 ± 7.33 | 24.38 ± 7.79 | 21.87 ± 2.97 | 0.426 |
| Water Intake (g/day),<br>Mean ± SD | 324.50 ± 89.20 | 356.80 ± 102.30 | 341.20 ± 95.60 | 0.452 |
| Food Intake (g/day), Mean ± SD | 94.60 ± 32.10 | 98.20 ± 28.70 | 96.80 ± 30.40 | 0.678 |
| Glossy Coat, n (%) | 5 (62.50%) | 21 (80.77%) | 13 (86.67%) | 0.294 |
| Normal Skin, n (%) | 7 (87.50%) | 22 (84.62%) | 14 (93.33%) | 0.824 |

NOTE: Continuous variables are presented as mean ± standard deviation; categorical variables are presented as number (percentage).
Abbreviations: HOM: homozygous (A31P/A31P); HET: heterozygous (A31P/N); WT: wild-type (N/N).
$P < 0.05$ was considered statistically significant.

### 3.2. Cardiac Structural and Functional Phenotypes

Based on the statistical analysis presented in Table 2 and Figure 2, significant differences in cardiac structural and functional parameters were observed among the three genotype groups. Structurally, the HOM group exhibited heterogeneous thickening of the left ventricular wall, with significantly greater thickness in the mid anteroseptal segment (HOM: 4.55 ± 0.83 mm vs. HET: 4.91 ± 0.53 mm vs. WT: 3.69 ± 0.54 mm, *P* = 0.018), the apical anterior segment (HOM: 4.01 ± 0.42 mm vs. HET: 3.53 ± 0.48 mm vs. WT: 3.96 ± 0.73 mm, *P* = 0.027), the apical lateral segment (HOM: 4.05 ± 0.51 mm vs. HET: 3.42 ± 0.54 mm vs. WT: 3.28 ± 0.45 mm, *P* = 0.016), and the mid inferior segment (HOM: 4.20 ± 1.05 mm vs. HET: 3.57 ± 0.38 mm vs. WT: 3.26 ± 0.61 mm, *P* = 0.013), compared to the other two groups. Concurrently, left atrial diameter (HOM: 15.79 ± 6.15 mm vs. HET: 11.63 ± 2.25 mm vs. WT: 11.29 ± 1.34 mm, *P* = 0.010) and left atrial volume index (HOM: 2.68 ± 1.96 ml/m² vs. HET: 2.39 ± 2.81 ml/m² vs. WT: 1.43 ± 0.29 ml/m², *P* = 0.024) were significantly larger in the HOM group.

**Figure 1.**
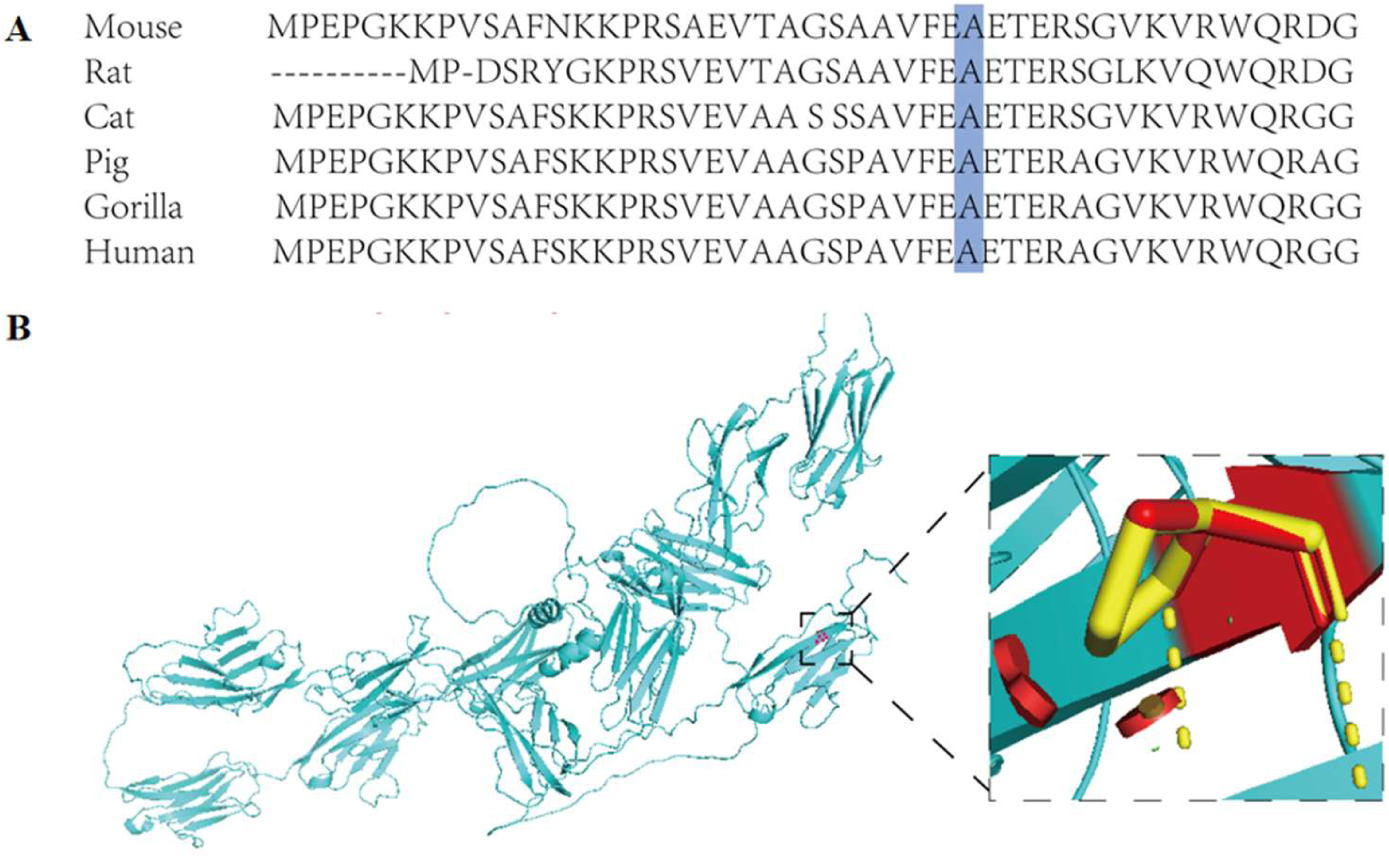
Functional prediction of protein encoding by MYBPC3-A31P mutation. (A) Analysis of sequence conservation. (B) The 3-dimensional structure of the amino acid p. Ala31Pro showing the conversion of valine for alanine.

**Figure 2.**
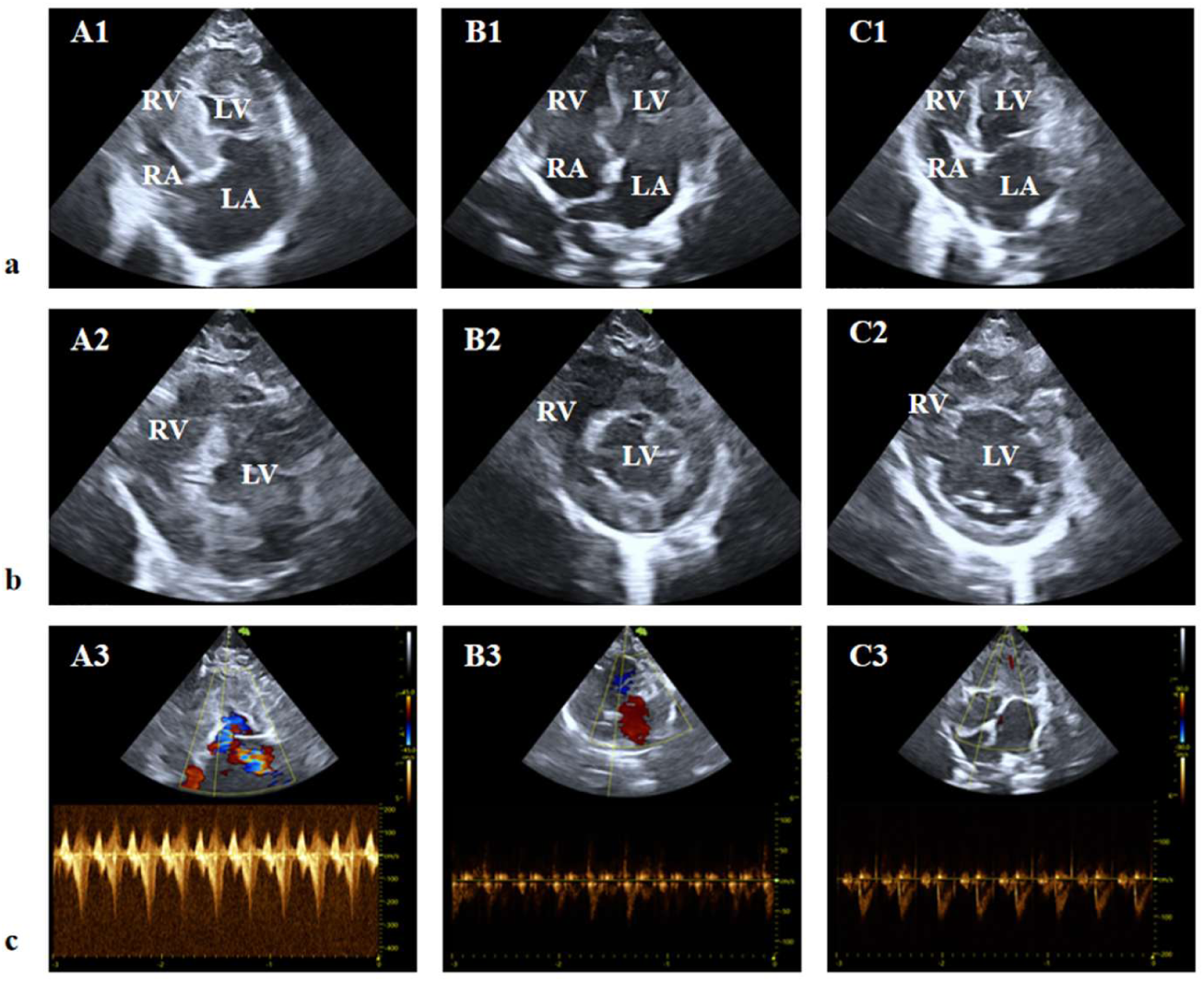
Examples of echocardiographic images. (a) Apical four-chamber view, (b) Left ventricular short-axis view (at the chordal level), (c) LVOT flow spectrum.

**Table 2.** Echocardiographic Parameters by MYBPC3 Genotype.

| Parameter | HOM<br>(A31P/A31P)<br>(n=8) | HET<br>(A31P/N)<br>(n=26) | WT<br>(N/N)<br>(n=15) | P-value |
| --- | --- | --- | --- | --- |
| LVOT PG (mmHg), Mean ± SD | 6.60± 10.30<sup>b</sup> | 3.46 ± 1.78 | 2.41 ± 1.43 | 0.030 |
| Basal inferoseptal(mm), Mean ± SD | 4.59±0.86 | 4.15±0.50 | 4.09±0.59 | 0.342 |
| Mid inferoseptal(mm), Mean ± SD | 4.37±0.57 | 4.00±0.67 | 4.11±0.65 | 0.301 |
| Apical septal(mm), Mean ± SD | 3.68 ± 0.36 | 3.57 ± 0.66 | 3.57 ± 0.73 | 0.828 |
| Basal anteroseptal(mm), Mean ± SD | 4.61 ± 0.91 | 4.06 ± 0.65 | 3.90 ± 0.54 | 0.180 |
| Mid anteroseptal(mm), Mean ± SD | 4.55±0.83<sup>b</sup> | 4.91±0.53 | 3.69±0.54 | 0.018 |
| Basal anterior(mm), Mean ± SD | 4.60 ± 0.76 | 4.06 ± 0.54 | 4.11± 0.71 | 0.196 |
| Mid anterior(mm), Mean ± SD | 4.33 ± 0.92<sup>b</sup> | 3.93 ± 0.53 | 3.86± 0.61 | 0.415 |
| Apical anterior(mm), Mean ± SD | 4.01 ± 0.42<sup>a</sup> | 3.53 ± 0.48 | 3.96± 0.73 | 0.027 |
| Basal lateral(mm), Mean ± SD | 4.33 ± 0.75 | 4.09 ± 0.59 | 3.85 ± 0.65 | 0.265 |
| Mid lateral(mm), Mean ± SD | 4.49 ± 1.00<sup>b</sup> | 4.03 ± 0.75 | 3.79 ± 0.60 | 0.138 |
| Apical lateral(mm), Mean ± SD | 4.05 ± 0.51<sup>a,b</sup> | 3.42 ± 0.54 | 3.28 ± 0.45 | 0.016 |
| Basal inferior(mm), Mean ± SD | 3.95 ± 0.86 | 3.77 ± 0.61 | 3.47 ± 0.50 | 0.312 |
| Mid inferior(mm), Mean ± SD | 4.09 ± 0.83 | 3.60 ± 0.60 | 3.50 ± 0.65 | 0.144 |
| Basal inferior(mm), Mean ± SD | 4.01 ± 1.37 | 3.63 ± 0.52 | 3.40 ± 0.51 | 0.376 |
| Mid inferior(mm), Mean ± SD | 4.20 ± 1.05<sup>b</sup> | 3.57 ± 0.38 | 3.26 ± 0.61 | 0.013 |
| Apical inferior(mm), Mean ± SD | 3.43 ± 0.53<sup>b</sup> | 3.13± 0.39 | 3.15 ± 0.53 | 0.217 |
| LVEF (%), Mean ± SD | 71.27 ± 6.00 | 71.65 ± 6.36 | 73.99 ± 5.91 | 0.537 |
| E wave (cm/s), Mean ± SD | 86.98 ± 23.71<sup>b</sup> | 70.34 ± 19.18 | 62.30 ± 10.50 | 0.033 |
| A wave (cm/s), Mean ± SD | 83.04 ± 23.70 | 69.58± 22.11 | 69.46 ± 17.87 | 0.447 |
| E/e' ratio, Mean ± SD | 9.60 ± 5.17 | 8.62 ± 2.54 | 8.30 ± 1.74 | 0.542 |
| SV (ml), Mean ± SD | 9.02 ± 2.27 | 9.12 ± 3.11 | 8.28 ± 2.02 | 0.626 |
| Left atrium diameter (mm), Mean ± SD | 15.79 ± 6.15<sup>a,b</sup> | 11.63 ± 2.25 | 11.29 ± 1.34 | 0.010 |
| LAVI (ml/m<sup>2</sup>), Mean ± SD | 2.68 ± 1.96<sup>b</sup> | 2.39 ± 2.81 | 1.43 ± 0.29 | 0.024 |
| LA:Ao Ratio, Mean ± SD | 1.50 ± 0.31<sup>a,b</sup> | 1.21 ± 0.28 | 1.29 ± 0.19 | 0.057 |

Abbreviations: LVOT, left ventricular outflow tract; LVEF, left ventricular ejection fraction; SV, stroke volume; LAD, left atrium diameter; LAVI, left atrial volume index. For remaining abbreviations, see Table 1.
NOTE: Continuous variables are presented as mean ± standard deviation.
<sup>a</sup> $P < 0.05$ versus group HET (A31P/N); <sup>b</sup> $P < 0.05$ versus group WT (N/N).

Functionally, the early diastolic mitral inflow velocity (E wave) was significantly elevated in the HOM group (86.98 ± 23.71 cm/s) compared to the HET (70.34 ± 19.18 cm/s) and WT (62.30 ± 10.50 cm/s) groups (*P* = 0.033), suggesting impaired diastolic function, although the E/e’ ratio showed no significant differences among the three groups (*P* = 0.542). Left ventricular ejection fraction and stroke volumedid not differ significantly across genotypes (all *P* > 0.05).

Regarding hemodynamic parameters, the LVOT-PG was notably higher in the HOM group (6.60 ± 10.30 mmHg) compared to the HET (3.46 ± 1.78 mmHg) and WT (2.41 ± 1.43 mmHg) groups (*P* = 0.030).

As shown in Figure 3, the presence of systolic anterior motion (SAM) of the mitral valve was exclusively observed in the homozygous mutant group. One cat (12.5%) in Group HOM was SAM-positive, whereas no cases were identified in the heterozygous or wild-type groups, and suggests that this phenotype may be related to gene dose.

**Figure 3.**
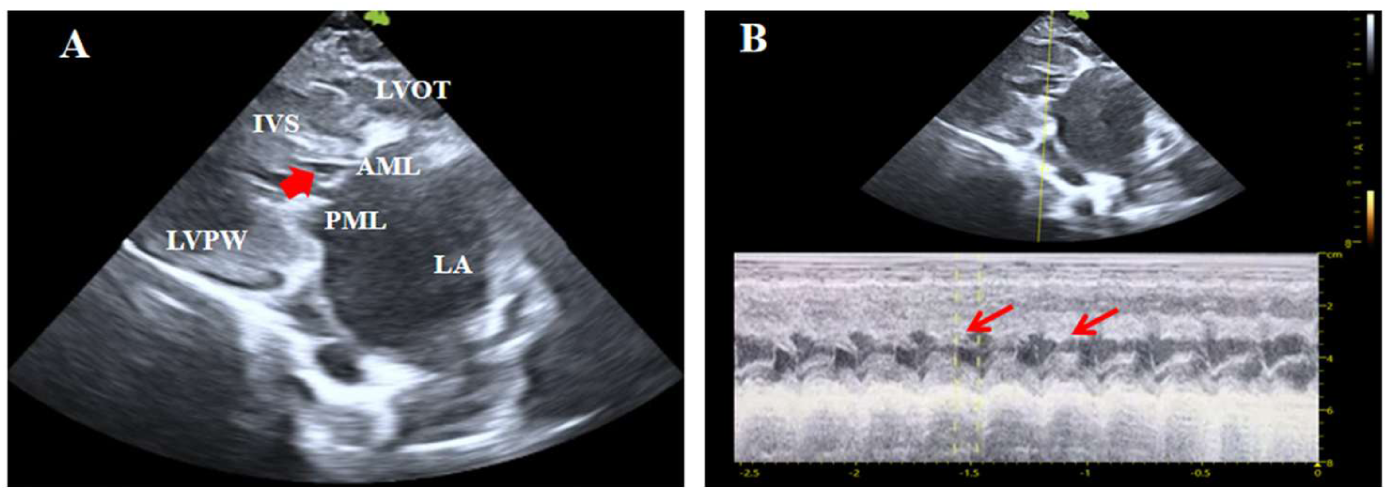
SAM of the mitral valve. (A) Parasternal long-axis view. Red arrow indicates anterior motion of the mitral valve (SAM). (B) M-mode echocardiogram. The small red arrow points to systolic anterior motion of the mitral valve leaflet; an abnormal upward deflection (arrowhead) is seen in the systolic CD segment.

Hematological and biochemical parameters are summarized in Table 3. Among all parameters evaluated, no significant differences were found among groups for the remaining parameters, including complete blood cell counts, serum amyloid A, albumin, total protein, alanine aminotransferase, hs-cTnT and myoglobin, possibly because the majority of our cats were at an early or subclinical stage, where such biomarkers are not yet elevated, and the sample size for certain genotypes may have limited power to detect small effect sizes^[11]^.

**Table 3.** Comparison of Key Hematological and Biochemical Parameters Among Genotype Groups.

| Parameter | HOM<br>(A31P/A31P)<br>(n=8) | HET<br>(A31P/N)<br>(n=26) | WT<br>(N/N)<br>(n=15) | *P*-value |
| --- | --- | --- | --- | --- |
| **Hematology** |  |  |  |  |
| WBC ($\times 10^9/L$) | 10.09±6.82 | 11.62±3.13 | 9.63±2.48 | 0.240 |
| Neutrophil Percentage (%) | 49.10±12.79 | 51.03±11.32 | 49.92±8.64 | 0.890 |
| RBC ($\times 10^{12}/L$) | 7.04±1.56 | 7.38±1.23 | 8.29±1.52 | 0.066 |
| HGB (g/L) | 129.70±21.55 | 130.30±16.21 | 141.00±22.21 | 0.196 |
| PLT ($\times 10^9/L$) | 249.00±94.81 | 254.80±86.33 | 249.40±122.40 | 0.981 |
| **Blood Biochemistry** |  |  |  |  |
| ALB(g/L) | 29.98±3.21 | 30.48±2.68 | 30.75±1.88 | 0.791 |
| TP(g/L) | 33.84±10.21 | 38.74±13.12 | 35.64±9.31 | 0.509 |
| Crea ($\mu\text{mol/L}$) | 119.40±54.92 | 108.20±16.20 | 122.50±14.57 | 0.205 |
| ALT (U/L) | 43.09±9.90 | 60.75±23.06 | 51.95±17.19 | 0.077 |
| **Myocardial Enzymes** |  |  |  |  |
| hs-cTnT(pg/mL) | 5.80±6.01 | 7.93±7.46 | 6.40±0.03 | 0.585 |
| Myoglobin(ng/mL) | 5.03±0.15 | 6.90±4.31 | 5.00±0.05 | 0.107 |

Abbreviations: WBC, white blood cell count; RBC, red blood cell; HGB, hemoglobin; PLT, platelet; ALB, albumin; TP, total protein; Crea, creatinine; ALT, alanine aminotransferase; hs-cTnT, high-sensitivity cardiac troponin T. For remaining abbreviations, see Table 1.
$P < 0.05$ was considered statistically significant.

### 3.3. Prediction of the Function of the Protein Encoded by the *MYBPC3*-A31P Mutation

To investigate the potential functional impact of HCM-associated genetic variation, we conducted an in-depth bioinformatic analysis of the *MYBPC3*-A31P missense mutation identified in Maine Coon cats. Using the Clustal Omega multiple sequence alignment tool available in the UniProt database, we performed a cross-species sequence conservation analysis of the mutation site. The results revealed that the alanine at position 31 (Ala31), which is altered by the mutation, is highly conserved across multiple mammalian species, including mouse, rat, pig, gorilla, and human (Figure 1A). The evolutionary conservation of this residue points to its potential functional significance, and mutations at this position may disrupt protein function, warranting further investigation. To further understand the mutation at the structural level, we used the PyMOL molecular visualization software to present three-dimensional structural models of *MYBPC3* before and after the mutation. In the rendered image (Figure 1B), the side chain of the wild-type alanine (Ala31) is marked in red, while the side chain of the mutant proline (Pro31) is highlighted in yellow.

## Discussion

This study systematically characterized the clinical phenotype of genotype-stratified Maine Coon cats carrying the *MYBPC3*-A31P mutation. The findings show a clear gene-dose effect: HOM cats had the most severe changes, HET cats fell in between, and WT cats remained unaffected. This gradient parallels the gene-dosage effects seen in human *MYBPC3*-associated HCM, supporting the model’s translational utility^[12]^. Baseline variables—including age, body weight, sex, heart rate, temperature, and routine blood/clinical parameters—did not differ across groups (all *P* > 0.05), confirming that the observed genotype-related differences were not confounded by these factors. Moreover, unlike many previous reports that enrolled HCM-affected cats from diverse breeding facilities or relied on clinically referred cases with heterogeneous backgrounds, all cats in the present study were raised under identical housing conditions and fed the same commercial diet. This uniformity minimizes the confounding effects of environmental variables—such as nutritional status, physical activity, stress levels, and management practices—on cardiac phenotype expression, further strengthening the attribution of the observed differences to the genotype rather than extrinsic factors.

Cross-species sequence conservation analysis revealed that alanine at position 31 (Ala31) of the *MYBPC3* protein is highly conserved across multiple mammalian species, including mouse, rat, pig, gorilla, and human. Highly conserved amino acid residues are typically located in functionally critical regions of proteins, and mutations in such regions are more likely to cause structural or functional abnormalities^[13]^. This finding aligns with the study by Van Dijk et al.^[13]^, which demonstrated using circular dichroism that the A31P mutation disrupts the conformation of the C0 domain of cMyBP-C, suggesting a “poison peptide” mechanism rather than haploinsufficiency. Three-dimensional structural modeling further suggests that the introduction of proline (Pro31) may alter local conformation, potentially affecting the interaction of the C0 domain with actin or the myosin regulatory light chain^[6]^. Such molecular-level alterations are also commonly observed in human HCM, where *MYBPC3* gene mutations represent one of the most frequent genetic causes^[14, 15]^.

Consistent with this molecular dysfunction, homozygous cats exhibited segmental left ventricular wall thickening—a pattern of asymmetric hypertrophy that is a hallmark of human HCM, with basal septal hypertrophy being most common^[16]^. Heterozygous cats did not show significant hypertrophy, indicating that at this age (mean 1.38 years) heterozygotes had not yet developed detectable morphological changes, in line with reports that homozygous cats manifest disease earlier than heterozygotes^[17]^.

Homozygous cats had significantly increased left atrial diameter and left atrial volume index (*P* < 0.05), with heterozygotes showing intermediate values. Left atrial enlargement is a morphological reflection of chronically elevated left ventricular filling pressure and diastolic dysfunction—parameters closely associated with disease severity in human HCM^[18]^. Studies in feline HCM have similarly confirmed that left atrial enlargement correlates with elevated left ventricular end-diastolic pressure and compensatory cardiac function^[19]^. The distribution of left atrial size observed in this study therefore suggests a relationship between left atrial enlargement and gene dose, reflecting differences in the degree of diastolic dysfunction across genotypes. Although the cross-sectional design precludes determination of the relationship with long-term outcomes, these findings provide important baseline data for future longitudinal follow-up. Systolic anterior motion of the mitral valve (SAM) was observed in only one homozygous cat (12.5%), with no cases identified in the heterozygous or wild-type groups. SAM is the primary mechanism of dynamic LVOT obstruction and the most common cause of heart murmurs in feline HCM^[20]^; in human patients, it is often associated with a more severe clinical course^[21]^. The observation of SAM exclusively in the homozygous group suggests that this phenotype may be related to gene dose, although the limited sample size necessitates validation in larger cohorts.

From a translational perspective, these findings have dual relevance: for veterinary practice, they highlight the need to avoid breeding combinations that yield homozygous cats, thereby reducing disease burden in the breed. For human HCM research, this well-characterized homozygous cohort provides a valuable platform for studying advanced disease mechanisms and evaluating therapeutic interventions. Compared to commonly used transgenic mouse models, this model offers distinct translational advantages. Engineered mouse models exhibit a rapid and homogeneous disease course, which often fails to replicate key clinical features of human HCM^[2]^. In contrast, cats in this study carry homologous *MYBPC3* mutations, and their disease progression closely resembles that of human HCM, providing an ideal platform for elucidating the association between genotype and phenotype.

## Limitations and Future Directions

This study also has several limitations. First, the relatively small sample size, particularly in the homozygous group (n = 8), limits statistical power and may have prevented detection of additional genotype-phenotype associations. Second, as a cross-sectional study, it cannot determine the temporal evolution of these phenotypes or their relationship with long-term outcomes. Further research is needed.

## Conclusion

In summary, this study provides comprehensive phenotypic characterization of *MYBPC3*-A31P-associated HCM in Maine Coon cats, demonstrating clear gene-dose effects on cardiac structure and function. Homozygous cats exhibit more severe hypertrophy, left atrial enlargement, and the exclusive occurrence of SAM, placing them at potentially higher risk for future clinical complications. These findings align with and extend previous observations on the role of gene dosage in feline HCM severity. The model holds high translational value due to its sharing of core pathological pathways with the human disease, thereby providing an efficient and reliable platform for future mechanistic exploration.

## Institutional Review Board Statement

This study was approved by the Institutional Animal Care and Use Committee of Yiji (Shanghai) Biotechnology Co., Ltd. (Approval No. YJ-2025-NA0015). All experimental procedures and animal welfare protocols complied with the NIH Guidelines for the Care and Use of Laboratory Animals.

## Acknowledgments

Author contributions: Conceptualization, Bo Wang, Jing Wang, Liwen Liu; methodology, Bo Wang; validation, Bo Wang, Xiaohe Shi, Ruoxuan Li; formal analysis, Xiaohe Shi, Ruoxuan Li; investigation, Bo Wang, Xiaohe Shi, Ruoxuan Li, Zixuan Yang; resources, Kai Liu; data curation, Xiaohe Shi, Yue Wang, Junzhe Huang; writing—original draft preparation, Xiaohe Shi; writing—review and editing, Xiaohe Shi, Bo Wang, Jing Wang; visualization, Xiaohe Shi, Ruoxuan Li, Zixuan Yang; supervision, Bo Wang, Jing Wang; project administration, Bo Wang, Jing Wang, Liwen Liu; funding acquisition, Bo Wang, Jing Wang, Liwen Liu. All authors have read and agreed to the published version of the article.

